# Evidence for strong purifying selection of human *47S* ribosomal RNA genes

**DOI:** 10.1101/2025.10.28.685169

**Authors:** Xufan Ma, Fiona Chow, Buz Galbraith, Daniel Sultanov, Eli K. Behar, Andreas Hochwagen

**Affiliations:** Department of Biology, New York University, New York, NY 10003, USA; Graduate Program in Data Science, New York University, New York, NY 10003, USA

**Keywords:** ribosomal rRNA genes, rDNA, conserved nucleotide elements, purifying selection

## Abstract

The multicopy *47S* ribosomal RNA (rRNA) genes are among the most highly expressed genes in the human genome, yet to-date essentially no disease-causing sequence variants have been identified. This lack of disease association is surprising, as defects in *47S* rRNA transcription and changes in ribosomal protein dosage, as well as nucleotide changes in the mitochondrial rRNA, all result in disease. The failure to identify rRNA-associated diseases may thus primarily stem from the experimental challenges associated with analyzing this chromosomally isolated high-copy gene family. Here, we used an evolutionary approach to test whether mutations in the human *47S* genes can have phenotypic consequences. By analyzing sequence variants among rRNA genes across >3,000 individuals from the high-coverage 1,000 Genomes Project, we demonstrate highly stratified variant abundance across the *47S* rRNA genes. In individual genomes, novel variants were frequently amplified in the transcribed spacer sequences and the evolutionarily young expansion segments, but rarely across the conserved *18S, 5*.*8S*, and *28S* rRNA-encoding sequences. Variant numbers and amplification were lowest in evolutionarily highly constrained nucleotide elements that are identical across >90% of sequenced eukaryotes. These results indicate that strong purifying selection acts to suppress copy number expansion of deleterious variants among the hundreds of *47S* rRNA copies and imply that deleterious variants in the *47S* rRNA have the potential to cause phenotypic consequences at very low copy numbers. As low-copy variant calls are rarely considered in association studies, this may explain why disease associations with *47S* rRNA variants have so far escaped detection.

**SIGNIFICANCE STATEMENT:** The rRNA genes are the most highly expressed genes in the human genome but there are almost no known diseases linked to sequence variants in the rRNA. We describe over 14,000 sequence variants that coexist within and between individuals and uncover signatures of strong purifying selection against deleterious variants. Our data indicate that deleterious rRNA variants cause sufficient fitness costs (and by extension, disease phenotypes) to be detected even against a massive backdrop of functional copies. As current disease-mapping algorithms generally ignore sequence variants that are only observed in a small percentage of sequencing reads, our data provide an obvious reason for the lack of disease associations.

## INTRODUCTION

Protein translation depends on the *28S, 18S, 5*.*8S*, and *5S* ribosomal RNAs (rRNAs), which form the catalytic core of the ribosome. To satisfy the high demand for ribosome production and protein synthesis, all four rRNAs are encoded by numerous gene copies. In the human genome, the small *5S* rRNA is encoded in a single highly repetitive locus on chromosome 1, whereas the *28S, 18S*, and *5*.*8S* rRNAs are produced from polycistronic *47S* transcripts that are encoded in repetitive ribosomal DNA (rDNA) arrays located on the short arms of the five acrocentric chromosomes, 13, 14, 15, 21, and 22 ^1-3^. An estimated 300-400 rRNA gene copies reside in these arrays, but copy number is highly variable among individuals and frequently changes from one generation to the next ^4-7^.

Although the *47S* rRNA genes are overall highly conserved, they harbor a substantial amount of sequence diversity. Most of this diversity maps to the large intergenic spacer sequences separating the *47S* transcription units and has been the subject of several recent studies ^8-12^ (**SI Appendix, Fig. S1a**). However, nucleotide variants have also been observed in the sequences coding for the *47S* transcript. Variation is particularly pronounced in the expansion segments ^10,13,14^, evolutionary young stem-loop structures with poorly defined functions that extend outward from the core of the ribosome ^15,16^. Variation has also been reported in the core RNA structures of the ribosome ^10,14,17^. Indeed, some of these variant rRNA sequences show differential expression among tissues as well as different epigenetic marks, implying tissue-specific regulation and possibly specialized functions for these variant ribosomes ^6,14,17-19^.

Recombination mechanisms, including gene conversion and unequal sister chromatid exchange, are highly active in the rDNA ^20,21^. In addition to changing total rRNA gene copy numbers, these mechanisms allow variants to increase or decrease in frequency among rDNA copies. As a result of these variant dynamics, sequence homogeneity among repeats is higher than expected from independent variant accumulation, leading to the concerted evolution of rDNA repeats ^22,23^. Most of the analysis of rRNA variants has focused on high-copy variants that are present in a large fraction of rRNA gene copies, with the expectation that high-copy variants are more likely to have functional consequences ^17,24^. Indeed, one recent analysis of the UK Biobank identified a significant correlation between variants in expansion segment 15L and body size traits ^24^.

What remains unclear is what fraction of all rRNA genes needs to harbor a particular variant before there are phenotypic consequences. Analyses of rDNA sequences in budding yeast (*S. cerevisiae*) indicate that this fraction may be quite low, as deleterious rRNA variants are strongly selected against, even if they are present at variant frequencies < 5% ^25^. Given the high degree of conservation of ribosomal RNAs, with blocks of near sequence identity across > 90% of sequenced eukaryotes ^26^, we predicted that stringent purifying selection also shapes the human rRNA sequences. To test this prediction, we took advantage of the relatively high sequence coverage offered by the most recent iteration of the 1,000 Genomes Project ^27^ to identify low-copy number nucleotide variants in the human rDNA at high confidence. This analysis yielded signatures of strong purifying selection across the *18S, 5*.*8S*, and large sections of the *28S* rRNAs.

## RESULTS

### Intra- and intergenomic sequence diversity of human *47S* sequences

To probe for signatures of purifying selection in the human rDNA, we focused on sequences of the *47S* transcript, which, unlike the intergenic spacers, exhibited largely uniform sequence coverage among samples of the 1,000 Genomes Project (**SI Appendix, Fig. S1a**). The primary *47S* transcript encodes the sequences of the *18S, 5*.*8S*, and *28S* rRNAs, as well as external and internal RNA spacer sequences (*5’ ETS, ITS1/2, 3’ ETS*) that are nucleolytically removed during ribosome biogenesis (schematic in **Fig. 1a; SI Appendix, Table S1**). These elements are predicted to be under varying levels of purifying selection ^25,28^. We identified variants by mapping sequencing reads to a prototype sequence of the *47S* gene. Comparing the average sequence coverage of the *47S* gene to the genome-wide sequence coverage yielded an estimated median rRNA gene copy number of 241 copies per haploid genome (min: 89, max: 544) for the samples in this dataset (**Fig. 1b**), in line with previous copy number measurements for the human genome ^4,5^.

**Figure 1.**
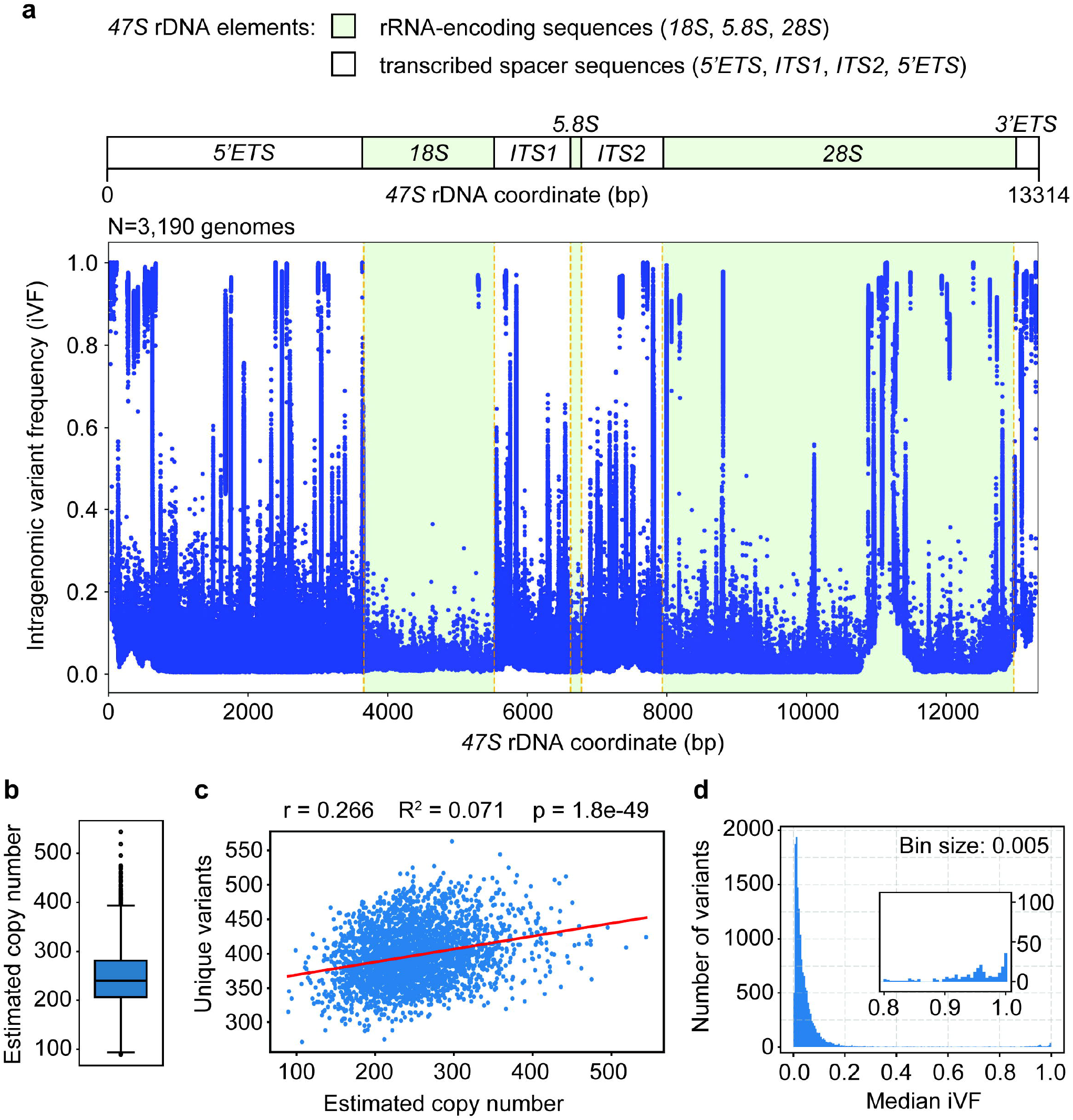
Distribution of sequence variants across the *47S* rDNA. (**a**) Top: schematic indicating the relative positions and lengths of individual *47S* elements. *ETS* - external transcribed spacer, *ITS* - internal transcribed spacer. rRNA-encoding sequences are shaded in pale green. Bottom: cumulative plot of all identified sequence variants and their associated frequencies observed in the high-coverage 1,000 Genomes Project data set. Each individual measurement is shown as a blue dot. Pale green shading and orange dashes indicate the positions and boundaries of the rRNA-encoding sequences, respectively. (**b**) Estimated *47S* copy number per haploid genome for each sample in the data set. Median is shown as black bar, boxes indicate 25-75% range, and error bars indicate 95% confidence interval. Measurements outside of this range are shown as individual dots to show overall spread of the data. (**c**) Scatter plot of the number of unique variants per sample as a function of estimated copy number. Each blue dot represents a sample, the regression line is shown in red with r, R^2^, and p values given. (**d**) Distribution of iVFs (binned into increments of 0.005) for all variants observed in the data set. Zoomed-in inset shows observed variant numbers at very high iVFs.

To identify variants in the *47S* gene, we adapted a pipeline previously used for rDNA variant analysis in budding yeast ^25^. This pipeline determines intragenomic variant frequencies (iVFs), i.e. the fraction of reads that code for a variant at each genomic position (Methods). Because of inherent challenges with sequencing long homopolymer runs, we excluded all variants mapping to homopolymers extending longer than 9 nucleotides (**SI Appendix, Table S2**). Benchmarking against the average 30-fold sequence coverage of the high-coverage dataset of the 1,000 Genomes Project indicated that the pipeline reliably and accurately recovered variants with iVFs ≥ 1% (**SI Appendix, Fig. S1b**). Below 1% iVFs, the fraction of false negatives increased substantially (∼75% false negatives at iVFs = 0.5%), although precision of recalled iVFs remained high (**SI Appendix, Fig. S1c**). The pipeline thus reliably recovers variants that are present in at least a handful of *47S* repeats.

Our pipeline also sought to exclude variants originating from likely pseudogenes. Most rDNA-derived pseudogenes have undergone sequence degradation ^29^ and thus did not enter our pipeline. As there is presently no evidence for rDNA-derived pseudogenes contributing to concerted evolution within the rDNA arrays, variants within pseudogenes are stably inherited and detectable as variants with consistently low iVFs across many samples ^30^. This characteristic rendered suspected pseudogenic variants readily identifiable at the population level. Based on these considerations, we excluded 290 presumed pseudogenic variants from further analysis (**SI Appendix, Dataset S1**). These variants were shared by at least 5% of the samples (n ≥ 160) in the dataset and showed consistent iVFs at very low frequencies, with little or no evidence of copy-number amplification across the whole dataset (see Methods). For a few of these variants, BLAST analysis identified perfect pseudogenic matches in the reference genome (GRCH38.p14). Most suspected pseudogenic variants were not identifiable in the reference genome and may map to the edges of the rDNA arrays, which harbor degenerating rDNA repeats that apparently escape recombination-mediated homogenization ^29^.

In total, we identified 14,878 unique *47S* rDNA variants across 3,190 samples of the 1,000 Genomes Project (**SI Appendix, Dataset S2**). These variants occurred throughout the *47S* (**Fig. 1a**), affecting more than 61.3% of the sequence (8,168/13,314bp). Approximately 75% of the variants were single-nucleotide variants (SNVs) and 25% were insertions or deletions (indels), ranging in size from 1-28 bp (median: 2 bp). This number is in line with analyses of short-read sequences ^10,17,18^, although a recent analysis using long reads suggested a substantially higher incidence of indels ^14^. As those novel indels occurred at very low iVFs, they may have been below the detection threshold of our analysis (**SI Appendix, Fig. S1b-c**). We observed a similar split between SNVs and indels for all rRNA-encoding sequences and spacer elements except for the 3’ external transcribed spacer (*3’ ETS*), which had a lower fraction of SNVs (**SI Appendix, Fig. S2a**). The number of unique variants correlated with estimated rDNA copy number (**Fig. 1c**), consistent with the notion that a larger number of rDNA repeats increases the opportunity for sequence variation.

Most variants occurred at iVFs below 0.2 (**Fig. 1d**), with a second peak at iVFs > 0.85 that is likely a consequence of rare variants in the prototype sequence (see below). Any given human genome harbored variants at 153 - 293 positions with various iVFs across all rDNA repeats (median: 214). In addition, some nucleotide positions displayed multiple non-reference variants (157 - 297 variants per genome; median: 218). Median iVFs per sample did not increase with estimated copy number (**SI Appendix, Fig. S2b**). Based on the high false-negative rate for very-low-frequency variants (**SI Appendix, Fig. S1b-c**), the true rDNA diversity within the 1,000 Genomes cohort is expected to be higher.

### Heritability of rDNA sequence variants

The vast majority of variants were either observed in a single individual or a small number of individuals, consistent with the strong overall conservation of the rDNA. Nevertheless, 61.4% of variants were observed in more than one sample (**Fig. 2a**), indicating either heritability or independent mutational events. The 1,000 Genomes Project comprises samples from 26 populations, each represented by 74 - 179 individuals ^27^. To get an estimate of the heritability of variants, we performed hypergeometric tests to compare the observed distribution of variants to the expected distribution based on the number of individuals in study populations in which a variant was identified. Variants were grouped into minor variants, i.e. variants observed in < 1% of samples (but in at least 2 samples), and medium-frequency variants, observed in 1-5% of samples. 3000 out of 6392 minor variants (46.9%) and 702 out of 731 medium-frequency variants (96%) showed significant enrichment in at least one study population. These data indicate that a substantial fraction of shared rDNA variants in this data set is the consequence of heritable events that propagate within these populations. These data also imply that the majority of *47S* variants called here are not the result of somatic mutations or sequencing artifacts.

**Figure 2:**
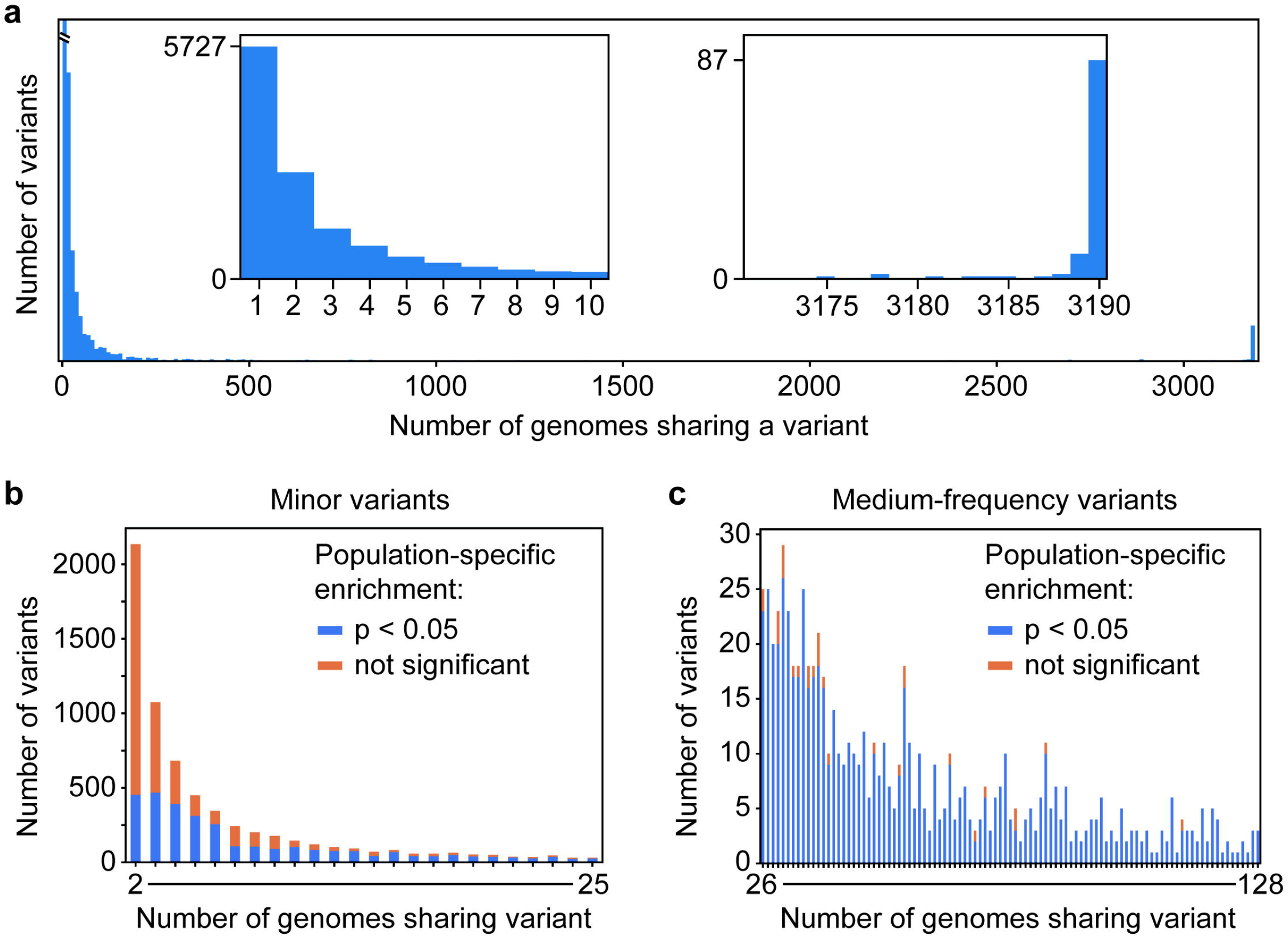
Heritability of *47S* variants. (**a**) Sharedness of variants among samples across the data set. Insets show zoom-ins at low and high levels of sharedness. (**b-c**) Analysis of population-specific enrichment of variants for (**b)** minor variants (observed in < 1% of samples, but at least 2 samples) and (**c**) medium-frequency variants (observed in 1-5% of samples). Bar graphs are sorted by the number of genomes sharing a given variant, with more common variants on the right. Blue color indicates significant population-specific enrichment (p < 0.05, hypergeometric test with Benjamini-Hochberg correction). Orange indicates no significant population-specific enrichment.

The sharedness analysis also revealed a small number of variants that are shared among > 99% of all samples (**Fig. 2a**). Some of these variants are likely the consequence of rare variants that were present in the rDNA prototype and thus reflect variants of a randomly sampled repeat that was chosen as reference. To test this possibility, we defined the major allele at every position within the dataset (see Methods) and used this information to construct a consensus prototype of the human *47S* rRNA region. Because of indel differences, the resulting consensus prototype is 13,365 bp long, and thus 51 bp longer than the original prototype. When we plotted the variant dataset against the consensus prototype, several common high-frequency variants disappeared (**SI Appendix, Fig. S3a-b**) and the number of variants shared by nearly all samples was reduced (**SI Appendix, Fig. S3c**), confirming that the high incidence of these variants was an artifact of rare variants present in the original prototype. The full sequence of the consensus *47S* prototype is given in **SI Appendix, File S1** and offers an alternative reference for future variant mapping experiments that is robust to reference-driven inflation of variant frequencies.

### Distribution of variants across the *47S* rDNA

Further analysis showed that variation was distributed unevenly across the *47S* rDNA. The fraction of variant nucleotides was lowest for the *18S* rRNA-encoding sequence as well as the *3’ ETS* (**Fig. 3a**). These two elements harbored the largest fraction of nucleotides that were identical across all reads in the dataset. In addition, variants with iVF > 10% were readily observed in the spacer sequences (5’ *ETS, ITS1/2*, 3’ *ETS*) but were notably depleted in the *18S* and *5*.*8S* rRNA-encoding sequences, as well as parts of the 28S rRNA-encoding sequences (**Fig. 1a, 3b**). By first approximation, unique variants represent individual mutational events in human history, although some variants likely arose independently more than once (see Discussion). By contrast, variant copy number reflects the amplification of a variant through the mechanisms of concerted evolution ^31^. Thus, to better capture the overall level variation, we calculated a variation score for each *47S* element by multiplying the fraction of variant nucleotides with the median level of iVFs observed across the dataset (see Methods). This analysis revealed consistently higher variation scores for the spacer regions compared to the rRNA-encoding sequences (**Fig. 3c-d**; t-test, p = 0.0064), strongly suggesting that rRNA-encoding sequences are under stronger purifying selection in the human population than the spacer regions.

**Figure 3:**
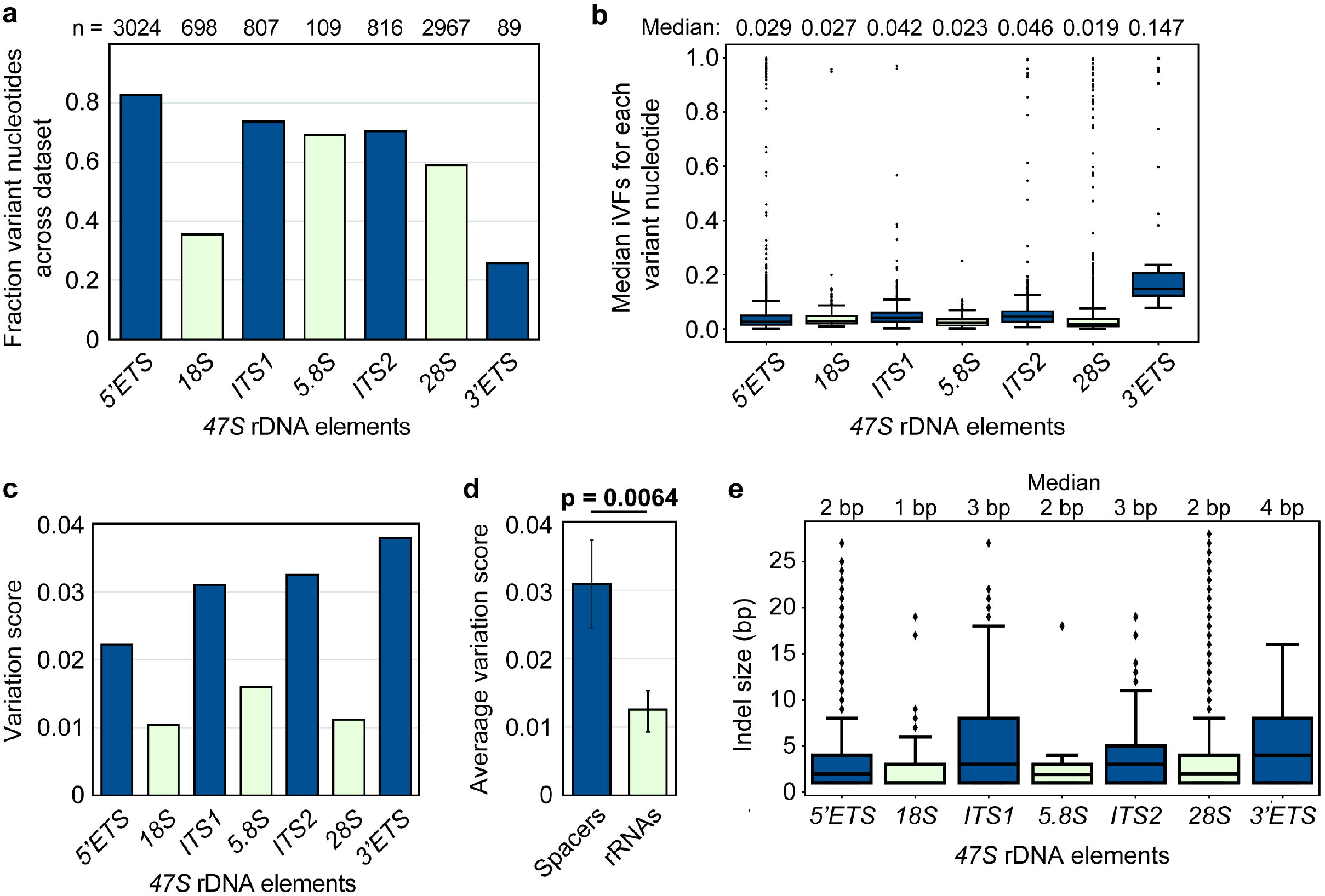
Differences in variant frequencies and identities across the *47S*. Analysis of variants as a function of *47S* elements in different *47S* elements. Spacer sequences are indicated in blue, rRNA-encoding sequences are shown in pale green. (**a**) Fraction of total nucleotides within each element for which variants were detected in the data set. (**b**) Box-and-whiskers plots showing the distribution of iVFs. Median is shown as black bar, boxes indicate 25-75% range and error bars indicate 95% confidence interval. Measurements outside of this range are shown as individual dots to show overall spread of the data. (c) Variation score calculated as the product of the fraction of variant nucleotides in (a) and the global median of the iVFs, noted on top of panel (b). (**d**) Averages of the variation scores of spacers and rRNA-encoding sequences shown in (c). Error bars indicate standard deviation. p = 0.0064; two-tailed t-test. (**e**) Box and whiskers plots showing the ranges and medians of indel sizes in the different *47S* elements.

We asked whether particular types of mutations are differentially enriched between spacer regions and rRNA-encoding sequences. Analysis of the 8,168 unique variant positions showed no consistent trends in the proportions of SNV and indels (**SI Appendix, Fig. S2a**) or in the relative proportion of transition and transversion mutations (**SI Appendix, Fig. S4a**) between spacer sequences and rRNA-encoding sequences. However, indel sizes were larger in spacer regions, suggesting more mutational freedom (**Fig. 3e**). In addition, G-C transversions were significantly more common in the spacers, while A-G transitions were more common in the rRNA sequences (**SI Appendix, Fig. S4b-e**). These mutational biases may be linked to the markedly higher G-C content of the spacers (**SI Appendix, Fig. S4f-g**) ^32^.

Overall, the number of unique variants (SNV and indels) correlated with the lengths of individual elements of the *47S*, both for rRNA-encoding sequences and for spacer sequences, but the number of variant positions was significantly depressed in the rRNAs compared to the spacer sequences (**SI Appendix, Fig. S5a**).

To further investigate the possibility of purifying selection in the rRNA-encoding sequences, we focused on the *28S* rRNA-encoding element. Although much of the *28S* rRNA that forms the core structure of the large ribosomal subunit is conserved, previous analyses have identified several elements of extreme conservation that map to the inner core of the large subunit (**Fig. 4a-b**, red). These conserved nucleotide elements (CNEs) comprise roughly 21% of the *28S* rRNA (**Fig. 4a**, and **SI Appendix, Table S3**) and are virtually identical across eukaryotes (> 90% sequence identity) ^26^. The extreme conservation across billions of years of evolution implies that any sequence change within the CNEs can be considered deleterious ^25^. By contrast, the expansion segments, variably sized, GC-rich stemloop structures that extend from the core of the large subunit (**Fig. 4a-b**, blue; **SI Appendix, Table S4**) and may have roles in ribosome regulation ^16,33^, have expanded notably in length during metazoan evolution ^15,34^. The expansion segments are often not completely resolvable by structure determination methods ^35^ and are hotspots of sequence variation in the human *28S* rDNA ^10^. Therefore, we classified the *28S* sequences into three classes: CNEs, expansion segments, and the rest of the *28S*.

**Figure 4:**
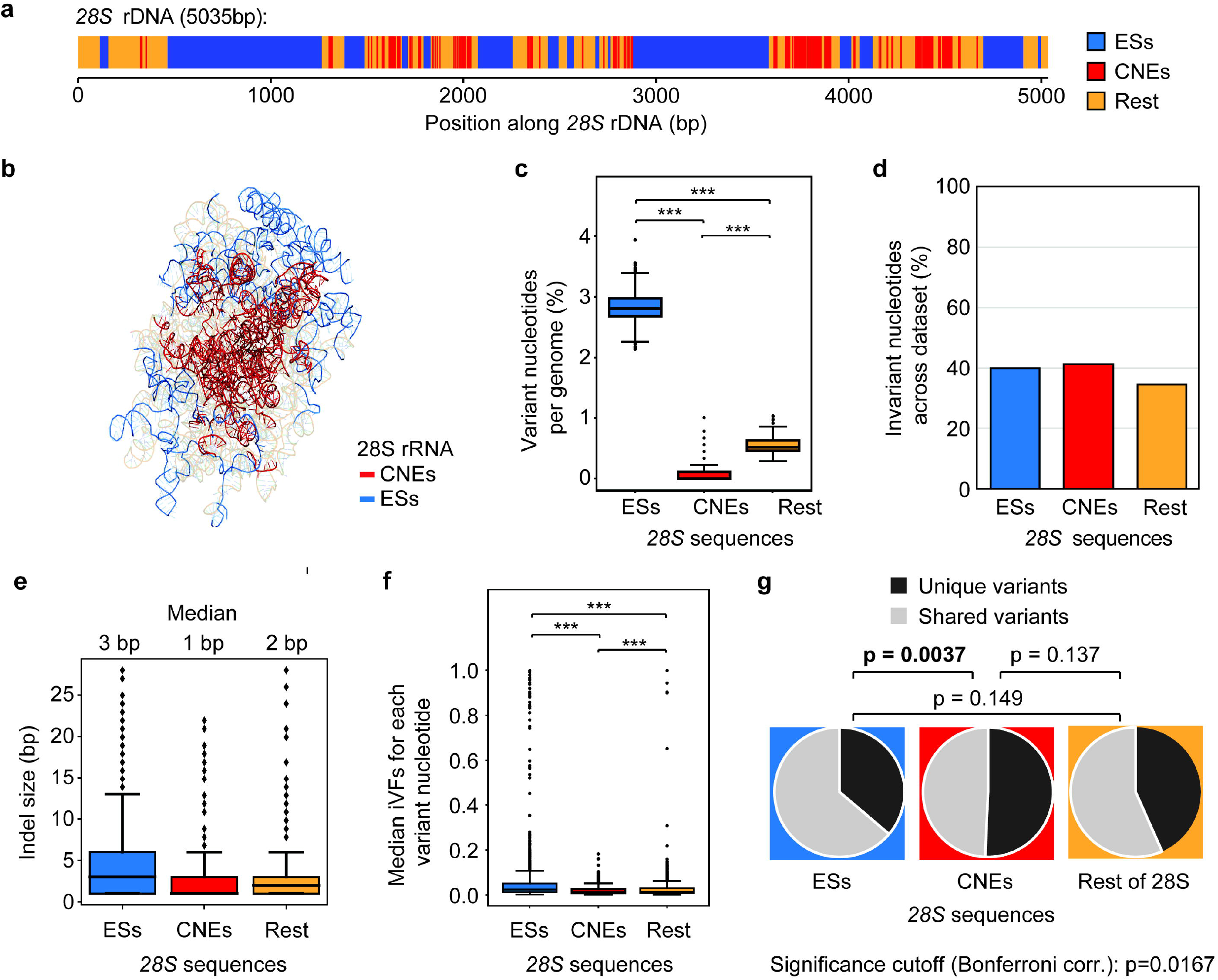
Evidence of purifying selection in the *28S* rDNA. Analysis of variants in the *28S* as a function of nucleotide elements. (**a**) Schematic of the positions of functional elements in the *28S* rDNA. Expansion segments (ESs) are indicated in blue, conserved nucleotide elements (CNEs) are shown red, all other *28S* sequences (rest) are shown in orange. (**b**) Structure of the human *28S* rRNA (PDB: 8QYX) indicating the location of CNEs in the core (red) and the well-ordered bases of the ESs at the periphery (blue). Most ES sequences do not have a uniform conformation and thus are not structurally resolved. (**c**) For each genome in the dataset, the percentage of nucleotides showing any level of variability in the *28S* elements was determined. The cumulative distribution of percentages among the 3190 genomes is shown as a box and whiskers plots. Median percentages are noted above the graph. (**d**) Percentage of nucleotide positions for which no variant was detected in the entire dataset. (**e**) Box and whiskers plots showing the ranges and medians of indel sizes. (**f**) For each identified variant, the median variant frequency was calculated across the dataset and plotted as a box and whiskers plot for all the variants in each nucleotide element. This approach was taken to avoid disproportionate effects of variants shared by multiple individuals on the distribution. (**g**) Pie charts indicating sharedness of *28S* variants among samples as function of nucleotide elements. Pairwise Chi-square analysis with Bonferroni correction.

Variant detection was strongly stratified among these three sequence classes. Individual genomes showed a median fraction of variant nucleotides (at any iVF) of 3.18% in the expansion segments, 0.11% in the CNEs and 0.51% in the rest of the *28S* (**Fig. 4c**). These differences are highly significant (two-way Kolmogorov-Smirnov test; p < 10^−5^). Importantly, the total fraction of nucleotides that were invariant across the entire dataset was similar for all three sequence classes (**Fig. 4d**), indicating similar mutation rates. These data suggest that purifying selection is strongest in the CNEs. In line with this interpretation, median indel sizes in the CNEs were significantly shorter than in the rest of the *28S* rRNA (**Fig. 4e**), and iVFs were lowest in the CNEs (**Fig. 4f**; two-way Kolmogorov-Smirnov test; p < 10^−5^). Finally, variants in the CNEs were also least likely to be shared among individuals (**Fig. 4g**). Taken together, these results imply that variants in the CNEs are deleterious and undergo purifying selection both at the level of variant amplification and at the level of inheritance. Our data also suggest that variants in the CNEs can cause these deleterious effects even when present at extremely low copy numbers.

## Discussion

Here we used comparative genomic analyses to reveal signatures of strong purifying selection across elements of the human *47S* rRNA genes. The significant depletion of high-copy variants in these segments suggests strongly dominant effects that can cause phenotypic consequences even against the backdrop of a large number of gene copies with the reference variant.

The elements of the *47S* rRNA genes exhibited differences in variation both in the frequency at which unique variants were detected and in the extent to which variants were amplified in copy number. By first approximation, unique variants reflect the mutational events across the data set. Assuming these events follow a Poisson distribution, we estimate that approximately 60% of such variants are the result of distinct mutational events and the remaining variants occurred independently two or more times in the study population. Supporting the notion that mutational events tend to be unique, we found that > 50% of variants shared between two individuals segregated in the same population group. However, there are clear limits to this model because the sequencing depth of the data set did not allow a comprehensive capture of variants with iVF < 1%. In addition, strong purifying selection may have suppressed the copy numbers of deleterious variants below our limit of detection (**SI Appendix, Fig. S5a**), thereby selectively reducing our ability to capture mutational events in the rRNA-encoding genes. It is also possible that mutation frequencies differ between spacer sequences and the rRNA-encoding sequences. This possibility is suggested by the notably higher GC content of the spacer sequences and expansion segments (**SI Appendix, Figs. S4f-g** and **S5b**), and the fact that differences in base composition can result in different mutation rates ^36,37^. However, we did not observe any obvious differences in the type of variants (transitions, transversions, indels) across the different elements of the *47S* that could hint at differential activity of repair mechanisms. Consistent with random mutational events, the number of unique variants correlated strictly with the lengths of the various spacers and rRNA-encoding segments. Uniform mutation accumulation is also in line with similar observations in the *S. cerevisiae* rDNA, which was sequenced at much higher coverage ^25,38^.

Our findings show that variants in the spacer regions and the expansion segments of the *28S* rDNA change readily in variant frequencies across individuals, whereas variant amplification was substantially more constrained in the conserved rRNA-encoding sequences. These data suggest that spacer regions are generally less mutationally constrained, either because only a subset of nucleotides in the spacers have important roles or because it is easier for cells to compensate for defects in ribosomal RNA processing than to cope with defective ribosomes.

The spreading of variants among rDNA repeats is fully consistent with the recombination mechanisms that drive the concerted evolution of rRNA genes ^22,23^ and shows that these mechanisms are highly active in the human population. Some of the recombination mechanisms, such as gene conversion, can mediate the transfer of single variants between repeats. However, we note that the mechanisms that underlie the frequent changes in rDNA copy numbers, such as unequal sister chromatid exchange or break induced recombination ^21^, amplify or delete the sequences of entire repeats and thus amplify or delete variants regardless of their location in the *47S*. The fact that high-frequency variants are nevertheless rarely observed in the structural RNAs of the ribosome (except the expansion segments) implies strong purifying selection that eliminates genomes in which deleterious variants have spread to too many rRNA gene copies.

The low frequency of variants in the structural RNAs is even more striking given that not all rRNA genes are expressed at any given time ^6,39^. Recent measurements of individual human rDNA arrays have revealed profound differences in epigenetic regulation and expression levels among arrays ^6,40,41^. Our data imply that silencing of rRNA genes must change (between tissues, between individuals, or over time), as permanent and heritable silencing of specific repeats would shield variants in those repeats from selection.

The fact that predicted deleterious variants rarely occur at variant frequencies higher than a few percent further indicates that these variants have the potential to poison cellular physiology even when most ribosomes do not carry this variant. Poisoning could be a result of the extreme cellular abundance of ribosomes and the fact that translation frequently involves multiple ribosomes on the same transcript. Whether these selective effects occur at the cellular or the organismal level remains to be determined. Regardless, these data predict that individuals who carry deleterious variants at elevated frequencies will likely experience deleterious phenotypes and disease and warrant a closer examination of predicted deleterious rDNA variants in disease association studies.

## Materials and Methods

### Read mapping and coverage

Using SRA Toolkit (v2.10.8), we obtained 3,190 high-coverage (30X) whole-genome sequencing (WGS) samples from the National Center of Biotechnology Information (Accessions: PRJEB37677 and PRJEB5507) ^27^. The downloaded SRA files were converted to paired-end FASTQ format and processed reads were aligned against a reference human rDNA prototype (GenBank: U13369.1) using Bowtie2 (v2.4.2). Although multiple rDNA sequences are available on GenBank ^9,11,42,43^, they are largely consistent across the *47S* gene.

### Variant calling

The resulting mapping data were sorted and compressed into .bam format and analyzed in a two-step variant calling process using LoFreq (v2.1.5) ^44^. First, alignment was adjusted around the insertions and deletions (indels) to minimize artifacts. Then the allele frequency of SNVs and indels was extracted across all reads and all positions, yielding an intragenomic variant frequency (iVF), i.e. the fraction of reads that code for a variant, at each genomic position of the rDNA prototype.

### Variant filtering

To limit potentially artifactual variant calls, we excluded homopolymer-associated variants and presumed pseudogenic variants from further analysis. As homopolymers are a common source of sequencing errors that would appear as reproducible variants in our analysis, we excluded variants that mapped to homopolymers. Homopolymers were defined as ≥ 10 consecutive identical nucleotides (C or G-rich), allowing 1 mismatch (**SI Appendix, Table S2**). In addition, we defined a list of 290 predicted pseudogenic variants (**SI Appendix, Dataset S1**), which were shared across at least 5% of individuals in the sample cohort (n ≥ 160) and displayed a concentrated low variant frequency profile that allows for a low level of outliers. Specifically, 97.5% (2 standard deviations from the mean) of variant frequencies of a predicted pseudogenic variant must fall under an iVF = 0.0714, which is a rounded estimate of 1/14, the minimum number of reads with an allele (1), divided by one of the lowest reported *47S* rRNA copy numbers in human (14) ^4^. This analysis was performed to account for the fact that a variant being physically present as a single copy in a genome can yield different iVFs depending on the total rDNA copy number.

### Benchmarking

To benchmark the sensitivity of our variant calling pipeline’s ability to capture variants at different levels of coverage, we generated synthetic samples using the NExt-generation sequencing Analysis Toolkit (NEAT) ^45^. The reads of one randomly selected subject’s genetic profile were down-sampled at different coverages, from 7x, 30x, 50x, 100x and 200x. Then synthetic variants were introduced at particular allele frequencies by mixing variant and reference reads in controlled ratios. The variant calling pipeline was benchmarked against these ground-truth datasets to obtain estimates of false-positive and false-negative errors as well as accuracy of iVF recall at each level of coverage.

### Copy-number estimates

To obtain a rough estimate of total *47S* copy number in a given sample, the average sequencing depth across the rRNA-encoding sequences of the *47S* was divided by the total sequencing depth of that sample (calculated by dividing total sequenced bases by the length of the reference genome GRCH38.p14). This calculation yielded an estimate of the average *47S* copy number per haploid genome. We chose to only include the rRNA-encoding sequences in this calculation because the spacer sequences have an unusually high GC content (**SI Appendix, Fig. S4g**). We note that the effects of high GC content on coverage and copy number estimates were overall mild (**SI Appendix, Fig. S1a** and **S1d**).

### Heritability analysis

To test for heritability, we tested for the enrichment of variants in specific populations. The underlying assumption is that study populations are genetically more related and thus heritable rDNA variants should be enriched in a given population. By contrast, if variants were primarily the results of independent mutational events (e.g. somatic mutations), then variants should not be linked to population. We opted to remove all children of other probands (n = 612) from this analysis, leaving 2,578 samples. Although coexistence of variants in parents and children offers evidence of heritability, we wanted to avoid confounding effects from the uneven distribution of parent-child pairs across the different study populations. To test for population-specific enrichment, we calculated the hypergeometric distribution comparing the observed distribution of any given variant in specific study populations with the null hypothesis that the variant should be observed proportional to the number of samples in the respective study populations (spanning between 74 and 179 samples). Results were corrected for multiple-hypothesis testing using the Benjamini-Hochberg correction.

### Consensus *47S* prototype

To define the consensus *47S* sequence of the 1,000 Genomes Project, we first established consensus *47S* sequences for each individual in the data set by defining the majority allele at each position of the prototype. We then defined the majority allele across the dataset by tallying the majority alleles among all individuals for a given position. These alleles were then concatenated to establish the consensus *47S* prototype (**SI Appendix, File S1**). Because of net difference of insertions over deletions among the majority alleles, the consensus *47S* prototype is 51 bp longer than the *47S* of U13369.1. This increase in length is the combined result of 38 indels compared to U13369.1.

### Analysis of *47S* and *28S* segments

The boundary coordinates of the elements of *47S* (using the U13369.1 reference) are given in **SI Appendix, Table S1**. The coordinates of the CNEs were defined by manual curation based on the sequences defined by ^26^ and are given in **SI Appendix, Table S3**. The coordinates of the expansion segments were taken from ^46^ and are given in **SI Appendix, Table S4**. The positions of CNEs and expansion segments were visualized on a cryo-EM derived structure of the *28S* (PDB: 8QYX; ^35^) using PyMOL.

### Calculation of variation scores

As described in the text, variation is observed in both the fraction of variant nucleotides (a proxy of incidence of mutational events) and the variation in iVFs (reflecting variant amplification events). Because of our inability to confidently detect variants with iVF < 0.01, these two forms of variation are not fully independent because strong iVF suppression by selection could result in iVFs that are undetectable, resulting in variants being erroneously counted as invariant. To account for this partial interdependence, we combined both forms of measurable variation by calculating a variation score that is the product of the fraction of variant nucleotides and the median level of iVFs observed across the dataset. We chose the median levels of iVFs for these calculations to avoid disproportionate effects of variants shared by multiple individuals.

### Estimating incidence of mutational events

In our analysis, 38.7% of nucleotides across the *47S* showed no detectable variants across the data set. Assuming a Poisson distribution, this incidence of invariant nucleotides yields an expectation λ = 0.94933. Based on this expectation, we estimate that 60% of the unique variants are the result of unique mutational events, whereas 40% of variants occurred independently 2 or more times.

## Author contributions

Conceptualization: X.M, D.S., A.H.; Investigation: X.M., F.C., B.G., A.H.; Code and formal analysis: X.M., F.C., B.G., D.S.; Illustration: E.B.; Writing - Original draft: X.M., D.S., A.H.; Writing - Review and Editing: X.M., F.C., B.G., E.K.B., D.S., A.H.

## Acknowledgements

This work was supported in part through the NYU IT High Performance Computing resources, services, and staff expertise. This work was supported by National Institutes of Health grant R35 GM148223 to A.H.

## Data availability

Coordinates of the variants identified in this study as well as all the coordinates used to define the different parts of the *47S* are given in **SI Appendix, Tables S1-S4** and **Datasets S1-S2**.

The computer scripts used for data analysis are available on Github: https://github.com/limaxf/Human_rDNA_project

