## Supplementary material for "Evidence for strong purifying selection of human *47S* ribosomal RNA genes": Figures S1-S5 and Tables S1-S4

**Supporting Information**

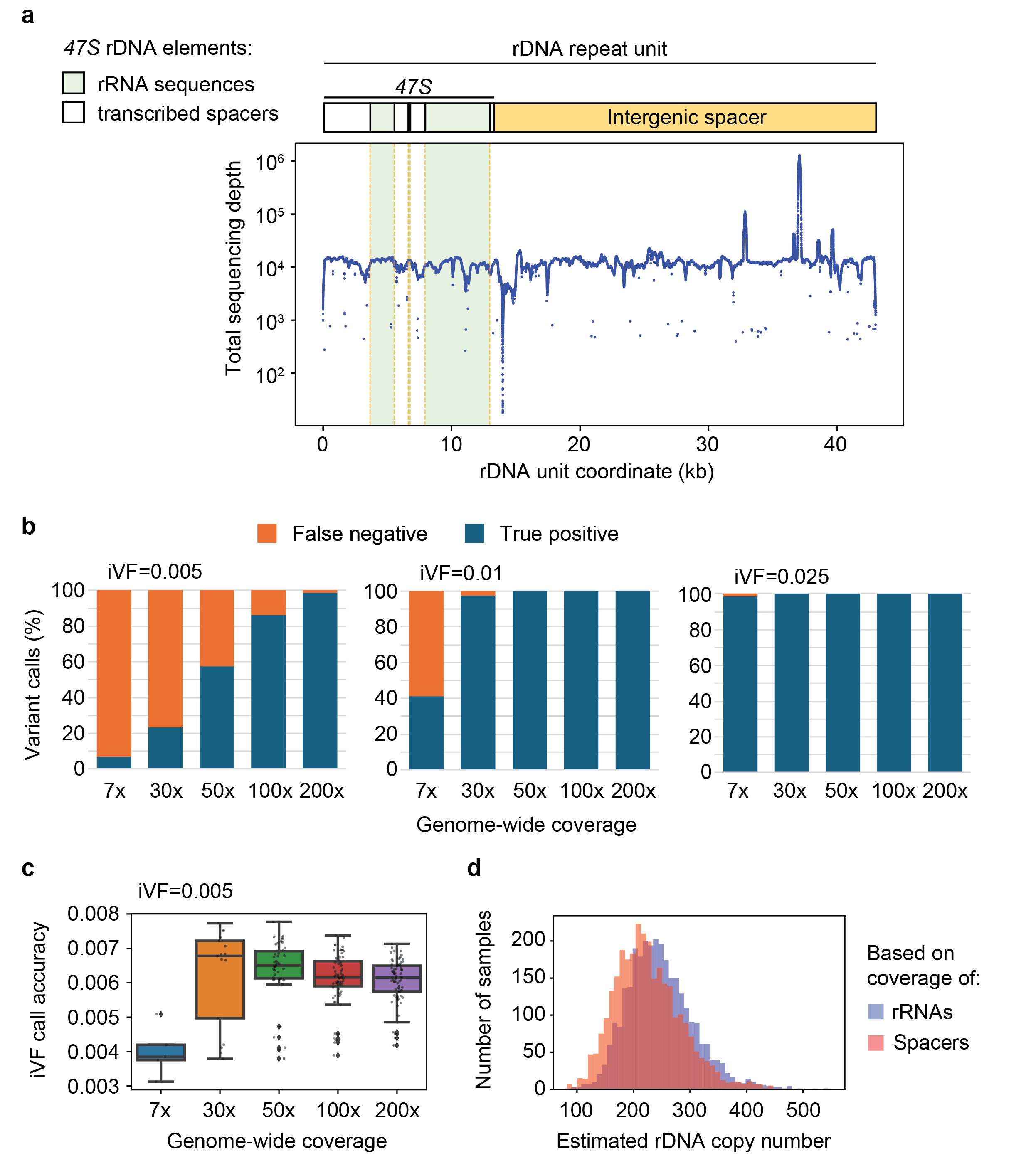

**Figure S1: Benchmarking of variant calling and copy number estimates.**

(**a**) Sequence coverage across an entire rDNA repeat. Top: schematic indicating the relative positions and lengths of the *47S* regions and the intergenic spacer (orange) that separates neighboring *47S*-encoding sequences. rRNA-encoding sequences are shaded in pale green, transcribed spacers of the *47S* are indicated in white. Bottom: sequence coverage across an rDNA repeat of a randomly selected genome from the high-coverage 1000 Genomes Project. Total sequencing depth is in the range of 5000-10000 because of the high copy number of the *47S* rDNA, but varies substantially in several sections of the intergenic spacer, likely because of local sequence composition or incompletely resolved repetitive sequences that also map outside the rDNA. (**b**) Variant calls recovered (true positives) and missed (false negatives) using a computationally spiked-in set of variants at different predetermined iVFs. iVFs used for benchmarking are shown in bold above the graph. Simulated genome-wide coverage is indicated below the graphs. The genomes in the high-coverage 1000 Genomes Project were sequenced to an average genome-wide coverage of 30x. (**c**) Box-and-whisker plots indicating precision of iVF calls at iVF = 0.005 as a function coverage. (**d**) Bar blot comparing rDNA copy-number estimates based on coverage of the rRNA-encoding sequences (purple) and transcribed spacer sequences of the *47S* (peach). The spacer sequences yielded lower estimates likely because their high GC content caused somewhat lower sequence yields.

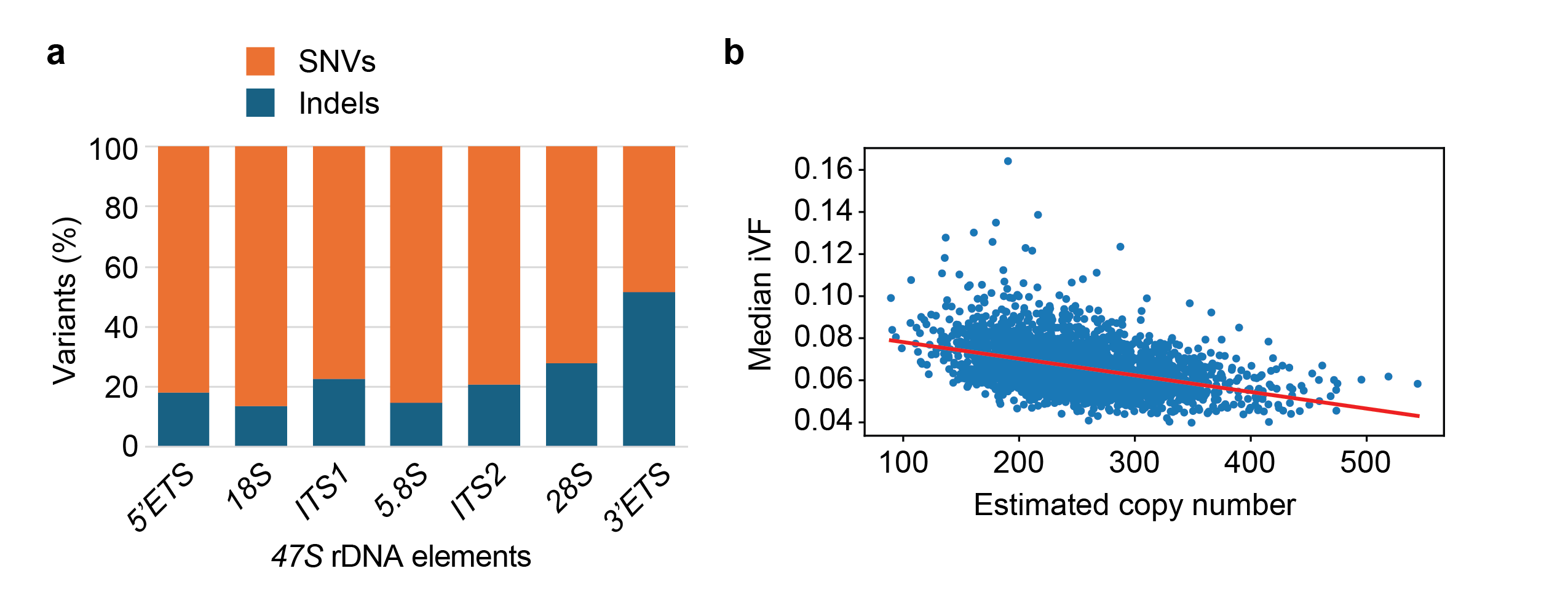

**Figure S2: Variants and variant frequencies across samples.**

(**a**) Relative occurrence of SNVs and indels among all detected variants is shown for the indicated elements. (**b**) Scatter plot of the median variant frequencies of each sample as a function of estimated copy number. Each blue dot represents a sample, the regression line is shown in red. The mild drop at higher copy-number estimates is related to the fact that higher copy numbers allow lower variant frequencies (i.e., the lowest possible iVF for a 100-copy sample is 0.01, whereas for a 500-copy sample it is 1/500 or 0.002).

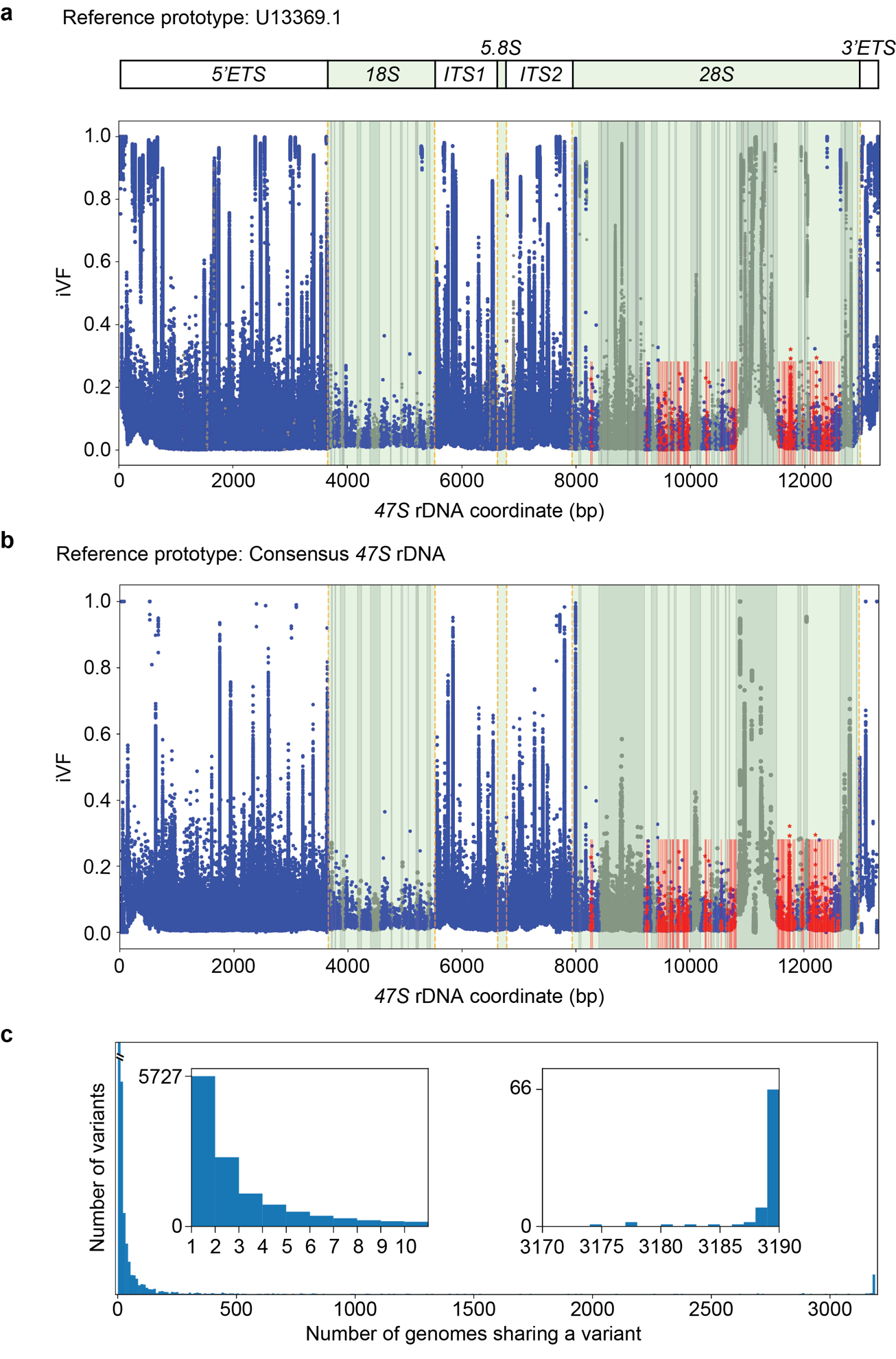

**Figure S3: Comparison of iVFs using the U13369.1 prototype or the consensus *47S* prototype.**

(**a**) Cumulative plot of all identified sequence variants and their associated frequencies observed in the high-coverage 1000 Genomes Project data set using U13369.1 as prototype. Each individual measurement is shown as a blue dot. Pale green shading and orange dashes indicate the positions and boundaries of the rRNA-encoding sequences, respectively. This panel is the same as in **Fig. 1a** but with CNEs indicated in red and other features, including ESs and homopolymers indicated in grey. The extensive peak in iVFs around position 11,000 is connected to the largest ES, *ES27* (length: 712 nt). (**b**) Cumulative plot of all identified sequence variants but corrected for majority alleles in the data set, i.e. using the consensus *47S* rDNA as the prototype. (**c**) Sharedness of variants among samples after correcting for majority alleles. Insets show zoom-ins at low and high levels of sharedness. The major effect compared to **Fig. 2a** is a loss of 21 highly shared variants that are unique to the U13369.1 prototype.

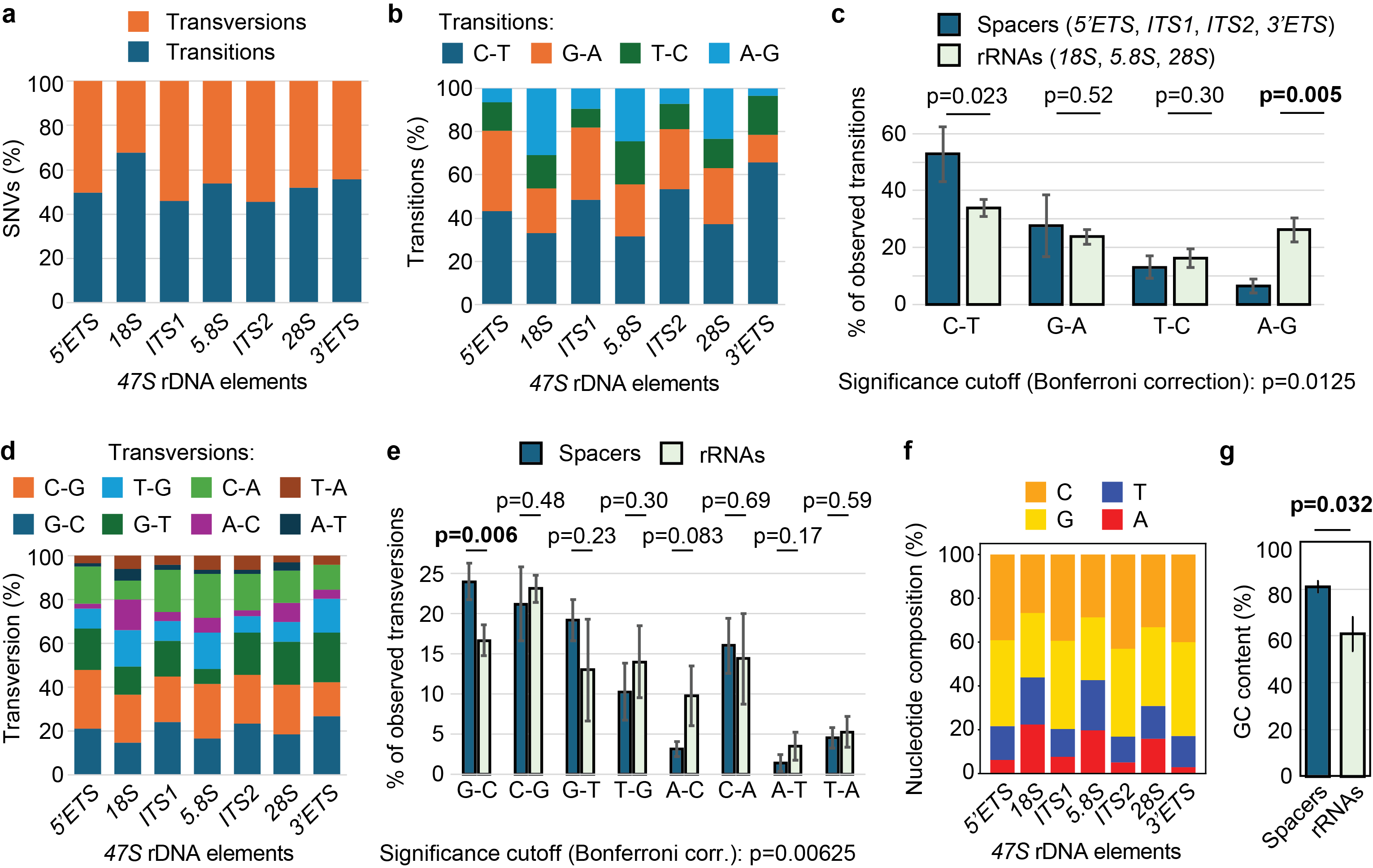

**Figure S4: Analysis of SNVs in different *47S* elements.**

Analysis of SNVs in different *47S* elements. (**a**) Relative occurrence of transitions and transversion in the indicated elements. (**b**) Breakdown of the individual types of transitions in the indicated elements. (**c**) Transition variants in spacers or rRNA-encoding sequences (e.g.: C-T indicates a transition from C to T). Error bars show standard deviation. P values were determined by Welch’s two-tailed t-test. P values below the Bonferroni-corrected significance level are highlighted in bold. (**d**) Breakdown of the individual types of transversions in the indicated elements. (**e**) Transversion variants in spacers or rRNA-encoding sequences (e.g.: C-G indicates a tranversion from C to G). Error bars show standard deviation. P values were determined by Welch’s two-tailed t-test. P values below the Bonferroni-corrected significance level are highlighted in bold. (**f**) Census of nucleotide frequencies in the indicated elements in the U13369.1 prototype. (**g**) Average GC content of spacer and rRNA-encoding sequences. Error bars show standard deviation. P values were determined by Welch’s two-tailed t-test.

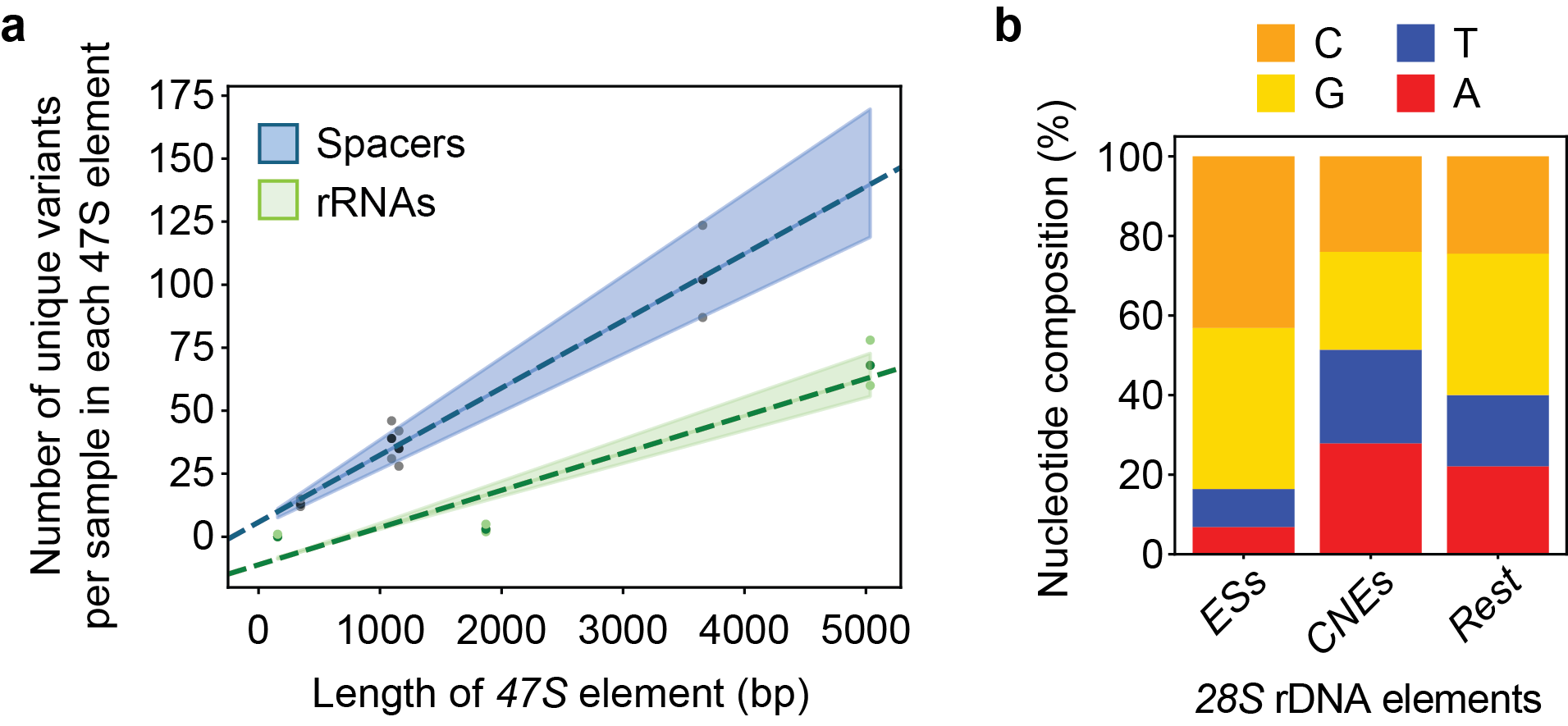

**Figure S5: Number of unique variants as a function of element length.**

(**a**) The average number of variants observed per sample in different *47S* elements is plotted as a function of the lengths of the different elements. Separate regression lines are shown for the spacer elements (black; r = 0.9942) and the rRNA-encoding elements (green; r = 0.9978). Shaded regions indicate the confidence intervals. The three data points are 5^th^ percentile, median (darker shade), and 95^th^ percentile. (**b**) Census of nucleotide frequencies in the indicated *28S* elements in the U13369.1 prototype.

**Table S1: Coordinates of 47S rDNA elements**

based on prototype (GenBank: U13369.1)

|  | **Start** | **End** | **Length** |
| --- | --- | --- | --- |
| *5'ETS* | 1 | 3656 | 3656 |
| *28S* | 3657 | 5527 | 1871 |
| *ITS1* | 5528 | 6622 | 1095 |
| *5.8S* | 6623 | 6779 | 157 |
| *ITS2* | 6780 | 7934 | 1155 |
| *28S* | 7935 | 12969 | 5035 |
| *3'ETS* | 12970 | 13314 | 345 |

**Table S2: Excluded homopolymers**

based on prototype (GenBank: U13369.1)

| **Start** | **End** | **Length** |
| --- | --- | --- |
| 3921 | 3930 | 10 |
| 8440 | 8449 | 10 |
| 8554 | 8563 | 10 |
| 8904 | 8913 | 10 |
| 9043 | 9052 | 10 |
| 9059 | 9068 | 10 |
| 9082 | 9092 | 11 |
| 10111 | 10129 | 19 |
| 10162 | 10171 | 10 |
| 10883 | 10894 | 12 |
| 10921 | 10930 | 10 |
| 11002 | 11012 | 11 |
| 11245 | 11261 | 17 |
| 11345 | 11354 | 10 |
| 11446 | 11455 | 10 |
| 11998 | 12007 | 10 |
| 12698 | 12711 | 14 |

**Table S3: Coordinates of conserved nucleotide elements in the *28S* rDNA**

based on prototype coordinates (GenBank: U13369.1)

| **Start** | **End** | **Length** |
| --- | --- | --- |
| 8256 | 8263 | 8 |
| 8284 | 8292 | 9 |
| 9234 | 9256 | 23 |
| 9439 | 9445 | 7 |
| 9454 | 9461 | 8 |
| 9483 | 9491 | 9 |
| 9511 | 9525 | 15 |
| 9546 | 9584 | 39 |
| 9589 | 9607 | 19 |
| 9654 | 9661 | 8 |
| 9663 | 9671 | 9 |
| 9698 | 9715 | 18 |
| 9772 | 9779 | 8 |
| 9784 | 9792 | 9 |
| 9795 | 9801 | 7 |
| 9810 | 9815 | 6 |
| 9882 | 9888 | 7 |
| 9890 | 9897 | 8 |
| 9899 | 9909 | 11 |
| 9912 | 9948 | 37 |
| 9952 | 9975 | 24 |
| 10266 | 10277 | 12 |
| 10278 | 10295 | 18 |
| 10327 | 10334 | 8 |
| 10549 | 10556 | 8 |
| 10659 | 10667 | 9 |
| 10710 | 10731 | 22 |
| 10735 | 10743 | 9 |
| 10766 | 10779 | 14 |
| 10784 | 10797 | 14 |
| 10803 | 10808 | 6 |
| 11543 | 11564 | 22 |
| 11602 | 11631 | 30 |
| 11640 | 11646 | 7 |
| 11648 | 11716 | 69 |
| 11718 | 11758 | 41 |
| 11759 | 11793 | 35 |
| 11807 | 11843 | 37 |
| 11958 | 11969 | 12 |
| 12089 | 12095 | 7 |
| 12097 | 12103 | 7 |
| 12130 | 12139 | 10 |
| 12155 | 12164 | 10 |
| 12202 | 12210 | 9 |
| 12279 | 12313 | 35 |
| 12315 | 12373 | 59 |
| 12398 | 12412 | 15 |
| 12417 | 12472 | 56 |
| 12502 | 12516 | 15 |
| 12597 | 12604 | 8 |

**Table S4: Coordinates of expansion segments in the *28S* rDNA**

based on prototype coordinates (GenBank: U13369.1)

| **Segment** | **Start** | **End** | **Length** |
| --- | --- | --- | --- |
| ESL5 | 8048 | 8090 | 43 |
| ESL7 | 8399 | 9199 | 801 |
| ESL9 | 9318 | 9421 | 104 |
| ESL10 | 9616 | 9645 | 30 |
| ESL12 | 9727 | 9763 | 37 |
| ESL15 | 10009 | 10190 | 182 |
| ESL19 | 10373 | 10428 | 56 |
| ESL20 | 10472 | 10509 | 38 |
| ESL24 | 10619 | 10639 | 21 |
| ESL26 | 10685 | 10698 | 14 |
| ESL27 | 10809 | 11520 | 712 |
| ESL30 | 11889 | 11947 | 59 |
| ESL31 | 11991 | 12058 | 68 |
| ESL39 | 12632 | 12839 | 208 |
| ESL41 | 12917 | 12930 | 14 |

**Provided as separate files:**

**Dataset S1: Excluded putative pseudogenic variants**

**Dataset S2: List of unique variants identified across 3,190 samples**

**File S1: Consensus *47S* rDNA prototype**
